# Microbiome features differentiating unsupervised-stratification based clusters of patients with abnormal glycometabolism

**DOI:** 10.1101/2022.12.19.521148

**Authors:** Ting Xu, Xuejiao Wang, Yu Chen, Hui Li, Liping Zhao, Xiaoying Ding, Chenhong Zhang

## Abstract

The alteration of gut microbiota structure plays a pivotal role in the pathogenesis of abnormal glycometabolism. However, the microbiome features identified in patient groups stratified solely based on glucose levels remain controversial among different studies.. In this study, we stratified 258 participants (discovery cohort) into three clusters according to an unsupervised method based on 16 clinical parameters involving the levels of blood glucose, insulin, and lipid. We found 67 cluster-specific microbiome features (i.e., amplicon sequence variants, ASVs) based on 16S rRNA gene V3-V4 region sequencing. Specifically, ASVs belonging to *Barnesville* and *Alistipes* were enriched in Cluster 1, in which participants had the lowest blood glucose levels, high insulin sensitivity, and high-fecal short-chain fatty acid concentration. ASVs belonging to *Prevotella copri* and *Ruminococcus gnavus* were enriched in Cluster 2, which was characterized by a moderate level of blood glucose, serious insulin resistance, and high levels of cholesterol and triglyceride. Cluster 3 was characterized by a high level of blood glucose and insulin deficiency, enriched with ASVs in *P. copri* and *Bacteroides vulgatus*. In addition, machine learning classifiers using the 67 cluster-specific ASVs were used to distinguish individuals in one cluster from those in the other two clusters both in discovery and testing cohorts (N = 83). Therefore, microbiome features identified based on the unsupervised stratification of patients with more inclusive clinical parameters may better reflect microbiota alterations associated with the progression of abnormal glycometabolism.

**IMPORTANCE:** The gut microbiota is altered in patients with type 2 diabetes (T2D) and prediabetes. The association of particular bacteria with T2D, however, varied among studies, which has made it challenging to develop precision medicine approaches for the prevention and alleviation of T2D. Blood glucose level is the only parameter in clustering patients when identifying the T2D-related bacteria in previous studies. This stratification ignores the fact that patients within the same blood glucose range differ in their insulin resistance and dyslipidemia, which also may be related to disordered gut microbiota. In addition to parameters of blood glucose levels, we also used additional parameters involving insulin and lipid levels to stratify participants into three clusters and further identified cluster-specific microbiome features. We further validated the association between these microbiome features and glycometabolism with an independent cohort. This work highlights the importance of stratification of patients with blood glucose, insulin, and lipid levels when identifying the microbiome features associated with the progression of abnormal glycometabolism.

## INTRODUCTION

A linkage between gut microbiota dysbiosis and type 2 diabetes (T2D) has been extensively explored (1–3). The alterations of the gut microbiota of patients with T2D are characterized by a decrease in the abundance of beneficial bacteria (e.g., *Akkermansia muciniphila*, the genera of *Bifidobacterium* and *Roseburia* spp.) and an increase in opportunistic pathogens including *Clostridium bolteae* and *Desulfovibrio* sp. (1, 4–6). The loss of potential butyrate-producing bacteria, such as *Faecalibacterium prausnitzii*, and the deficiency in butyrate production are the most common findings in patients with T2D as well as in patients with prediabetes (1, 3, 5–7). Nonetheless, the association of particular bacteria with T2D has varied among studies. For instance, Zhang *et al*. found that *A. muciniphila* had a lower abundance whereas *Clostridiales* sp. SS3/4 had a higher abundance in patients with T2D (4), but the abundances of these two taxa showed the opposite result from those reported in the study by Qin (1). This confusion has led to difficulty in using bacterial features to elucidate the mechanism of gut microbiota in the development of T2D for developing new biomarkers and therapies.

One possible reason for the challenge in identifying gut microbial characteristics in patients with T2D and prediabetes is that all of the previous studies have solely used blood glucose levels as the criteria to stratify their cohorts. Clinical investigation, however, has shown that people within the same blood glucose range as defined by the American Diabetes Association (ADA) and World Health Organization (WHO) criteria, are heterogeneous in insulin sensitivity and islet β-cell function (8, 9). Insulin resistance and islet β-cell dysfunction are the two pathogenesis of abnormal glycometabolism and occur much early than the onset of hyperglycemia (10). In addition, the condition of dyslipidemia, a risk factor of T2D and diabetic complications, is varied in people within the same blood glucose range. For example, in the Framingham Heart Study, 19% of the men and 17% of the women with diabetes had increased total plasma triglyceride levels, and the prevalence of increased total plasma triglyceride levels in men and women with normal blood glucose levels was 9% and 8% (11). Recently, apart from blood glucose levels, additional variables related to insulin resistance and sensitivity as well as dyslipidemia have been proposed for use in glycometabolism classifications (12–14). In addition to blood glucose levels, a study classified individuals, including healthy people and people with prediabetes, into six distinct clusters by including parameters such as insulin secretion, insulin resistance, and high-density lipoprotein-cholesterol (HDL), and followed them for 4.1 years and 16.3 years (in two cohorts, respectively) (12). Longitudinal follow-up revealed that different clusters had different risks of diabetes and diabetic complications. Individuals in one of the clusters experienced slower progression to overt T2D than those in other clusters but had a higher risk of nephropathy. In addition to the hyperglycemia, the disorder of gut microbiota has been shown to have a causative role in the progression of obesity and insulin resistance both in mouse models and human studies by gut microbiota transplantation. For example, a strain *Enterobacter cloacae* B29 isolated from a morbidly obese human’s gut induced obesity and insulin resistance in germfree mice (15). A gut microbiota transplantation study on humans revealed that gut microbiota from lean donors increased the insulin sensitivity in obese recipients with metabolic syndrome (16) Additionally, the gut microbiota could regulate insulin resistance by producing metabolites, such as imidazole propionate, which could directly impair insulin signaling at the level of insulin receptor substrate (17). Furthermore, a study in germ-free and conventionally raised mice showed that the gut microbiota had an effect on the host’s serum lipidomes, especially the triglyceride levels (18). In obese mice fed a high-fat diet, supplementation with probiotics, such as *Lactobacillus curvatus* alone or together with *L. plantarum*, reduced cholesterol in plasma; supplementation with *Bifidobacterium* spp. decreased the levels of circulating triglycerides and low-density lipoprotein-cholesterol (LDL), and increased the HDL level (19–21). Therefore, it is essential to consider insulin and lipid levels in the criteria for cohort stratification when studying the association of glycometabolism and gut microbiota.

In this study, to identify the microbiome features that correlate with progression of abnormal glycometabolism, we classified 258 individuals into three glycometabolism clusters according to variables related to blood glucose level, insulin resistance, and dyslipidemia. We found that the gut microbiota structure was different among the three glycometabolism clusters, and the characteristics of gut microbiota were associated with metabolic phenotypes. Moreover, we identified cluster-specific microbiome features and further validated their association with glycometabolism with a testing cohort.

## RESULTS

### **Unsupervised**-stratification based **clusters of patients with abnormal glycometabolism**

During a survey of patients with T2D conducted at Sijing Community Health Service Center of Shanghai Songjiang District, we recruited 267 participants as a discovery cohort and examined 33 bioclinical parameters of these participants. According to ADA criteria (8), 84 participants had normal glucose tolerance (NGT), 52 had isolated impaired fasting glucose (IFG), 34 had isolated impaired glucose tolerance (IGT), 38 had combined glucose intolerance (CGI), and 59 had T2D. We observed that the coefficients of variation (CV) of insulin levels for the oral glucose tolerance test (OGTT), homeostatic model assessment for insulin resistance (HOMA-IR), β-cell function (HOMA-β), serum triglyceride level, and cholesterol levels (total cholesterol, HDL, LDL) were high in participants with the same classification based on ADA criteria (Fig. 1A). We evaluated insulin resistance and insulin secretion using the HOMA-IR and HOMA-β index, which also showed significant variations in participants within the same glucose range (Fig. 1B, Fig. S1). These results confirmed that the subjects with same glucose level had high heterogeneity of metabolic state.

**FIG 1.**
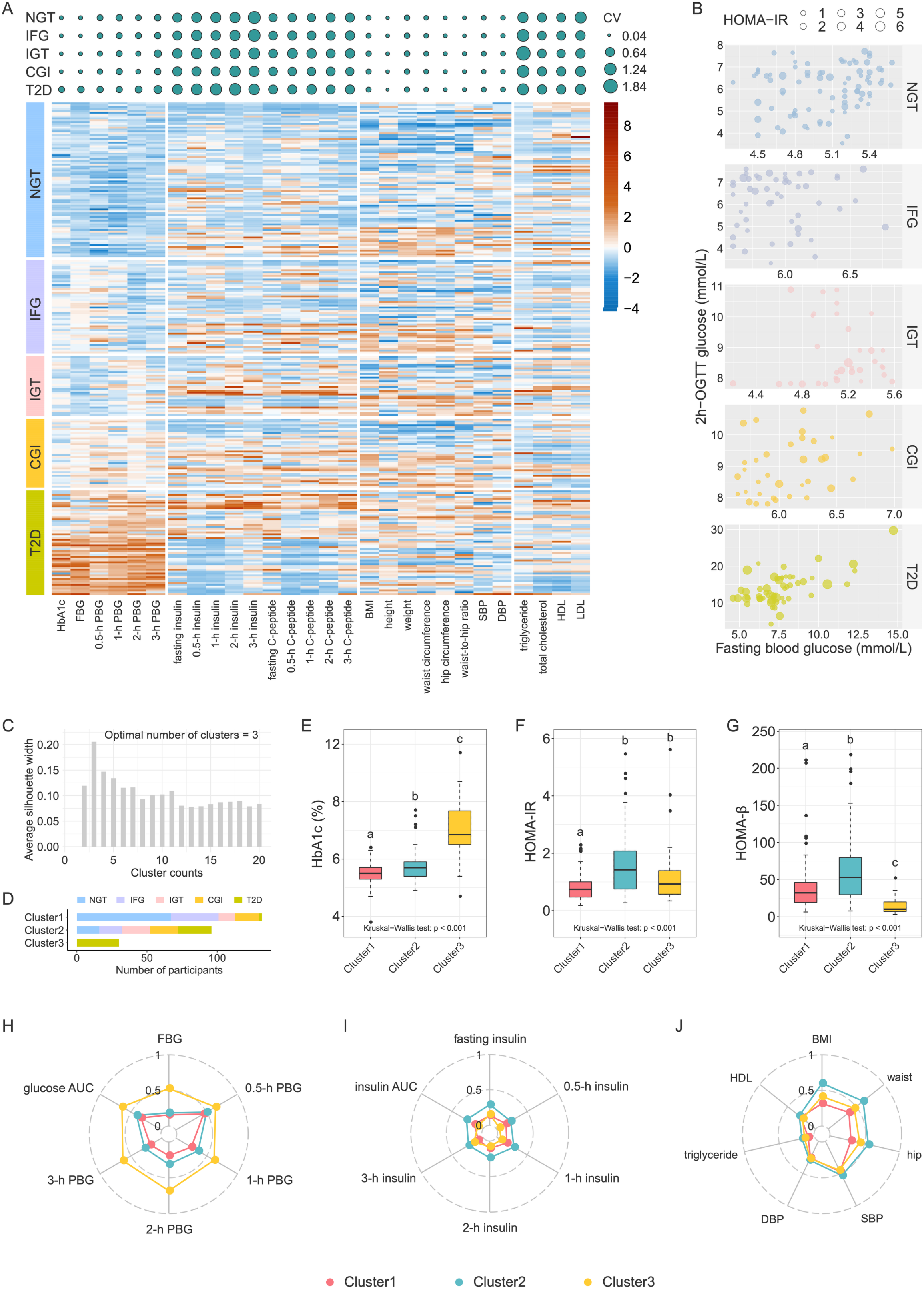
Metabolic characteristics of the unsupervised-stratification based clusters. **(A)** Top panel: coefficient of variation (CV) of clinical variables in each ADA group. The size of the circle indicates the CV value. Bottom panel: Heatmap of clinical variables. The values were scale-transformed by column. **(B)** Variations of HOMA-IR among members within the same blood glucose range. **(C)** Silhouette coefficient corresponding to the number of clusters from 2 to 20. **(D)** The number of participants in each cluster, with colors indicating glycemic categories (NGT, normal glucose tolerance; IFG, impaired fasting glycemia; IGT, impaired glucose tolerance; CGI, combined impaired fasting glycemia and impaired glucose tolerance; T2D, type 2 diabetes). Comparisons of **(E)** HbA1c, **(F)** HOMA-IR, and **(G)** HOMA-β among clusters. Boxes show the medians and the interquartile ranges (IQRs), the whiskers denote the lowest and highest values that were within 1.5 times the IQR from the first and third quartiles, and the outliers are shown as individual points. Kruskal-Wallis test *P*-value is shown at the bottom of each plot. The Wilcoxon rank-sum test was used for comparisons between two clusters (adjusted by FDR). Clusters with common characters were not significantly different (FDR > 0.05). The radar chart shows the median values of clinical parameters related to **(H)** blood glucose levels, **(I)** blood insulin levels, and **(J)** lipometabolism. Each spoke in the chart represents one cluster.

Then we reclassified the same group of participants in the discovery cohort according to variables including HbA1c, OGTT-derived glucose levels (five-time-point blood glucose levels during OGTT) and insulin levels (five-time-point blood insulin levels during OGTT), anthropometric variables (body mass index [BMI], waist circumference, hip circumference), as well as variables related to dyslipidemia and insulin resistance (fasting triglyceride, HDL cholesterol). We standardized the variables before the clustering procedure. We excluded nine participants (NGT = 1, IFG = 2, IGT = 2, CGI = 1, T2D = 3) with at least one outlier variable. By using the K-Mediods clustering algorithm (22), we classified 258 participants into three clusters that were determined according to the maximum silhouette coefficient (silhouette coefficient = 0.21) (Fig. 1C and 1D). The participants with NGT and prediabetes were clustered only into Clusters 1 and 2. Specifically, 80.7% participants with NGT were clustered into Cluster 1 and the rest with NGT were clustered into Cluster 2. In addition, 68% with IFG, 37.5% with IGT, and 45.9% with CGI were clustered into Cluster 1 and the remainder were clustered into Cluster 2. In contrast, most of T2D (53.6%) were clustered into Cluster 3, only 3.6% and 42.9% were clustered into Clusters 1 and 2, respectively. Finally, the Jaccard coefficient means for the three clusters were greater than 0.7 (0.78, 0.71, and 0.83, respectively), which indicated the three clusters were stable.

Compared with Cluster 1, Clusters 2 and 3 showed more serious disruptions in glycometabolism and lipometabolism and an increased inflammatory state (Table 1). The levels of HbA1c, FBG (fasting blood glucose), 0.5-h PBG (postprandial blood glucose), 1-h PBG, 2-h PBG, and 3-h PBG were lowest in Cluster 1 and highest in Cluster 3 (Fig. 1E and 1H, Table 1). The insulin levels during OGTT except 3-h insulin level were significantly higher in Cluster 2 than in Clusters 1 and 3 (Fig. 1I, Table 1), and the HOMA-IR index was significantly higher in Clusters 2 and 3 than in Cluster 1 (Fig. 1F), indicating that Cluster 1 had the highest insulin sensitivity, and Cluster 2 had compensatory secretion insulin, whereas the Cluster 3 was insulin deficiency (Fig. 1G). In addition, BMI, waist-to-hip ratio (WHR), and systolic blood pressure (SBP) were significantly higher in Cluster 2 than those in the other two clusters (Fig. 1J, Table 1). The levels of total cholesterol, triglyceride, and leptin were significantly higher in Clusters 2 and 3 than those in Cluster 1 (Fig. 1J, Table 1). The lipopolysaccharide (LPS)-binding protein (LBP) level, an indicator of chronic inflammation, was lowest in Cluster 1 and highest in Cluster 3 (Table 1).

**Table 1.**
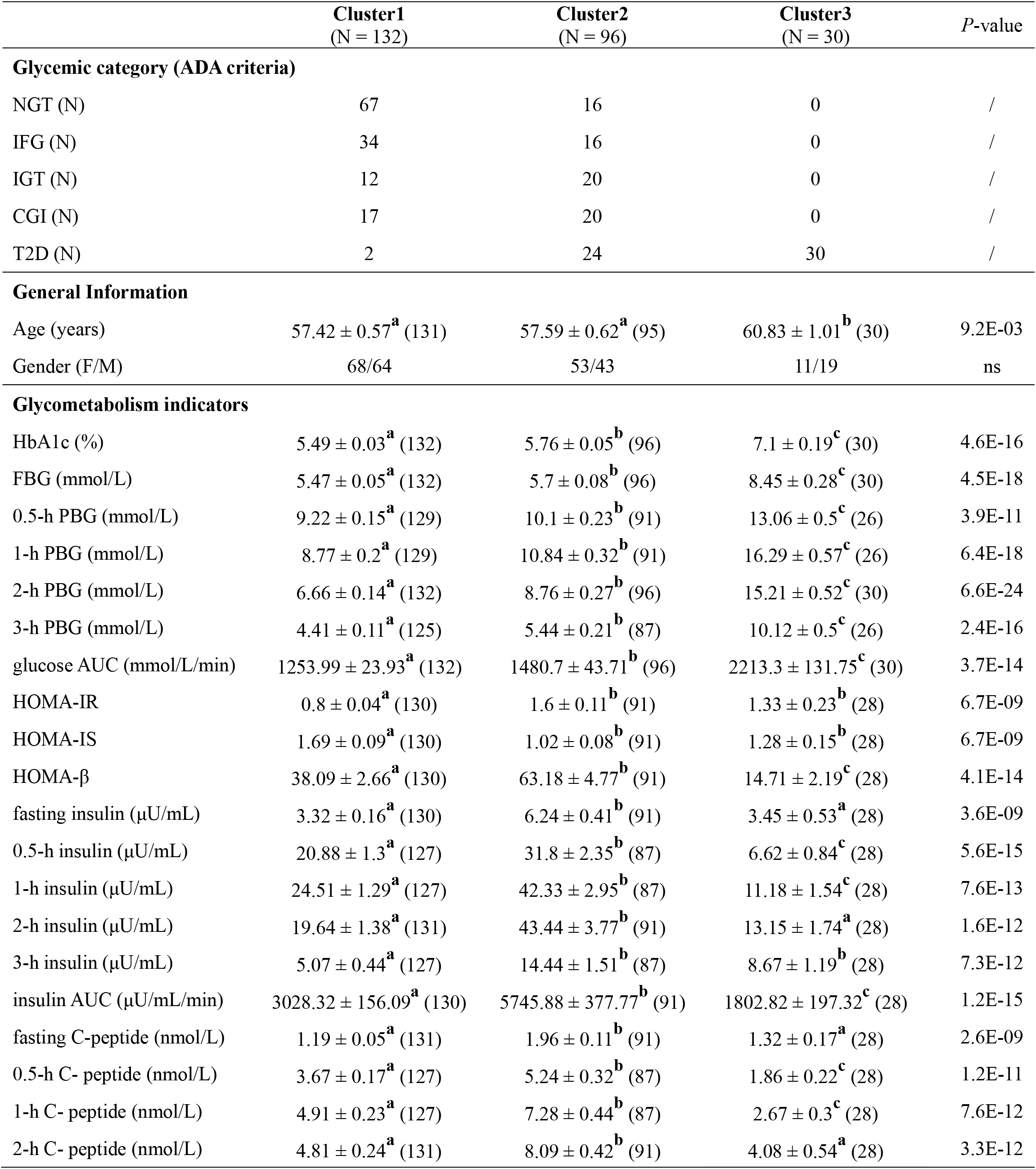
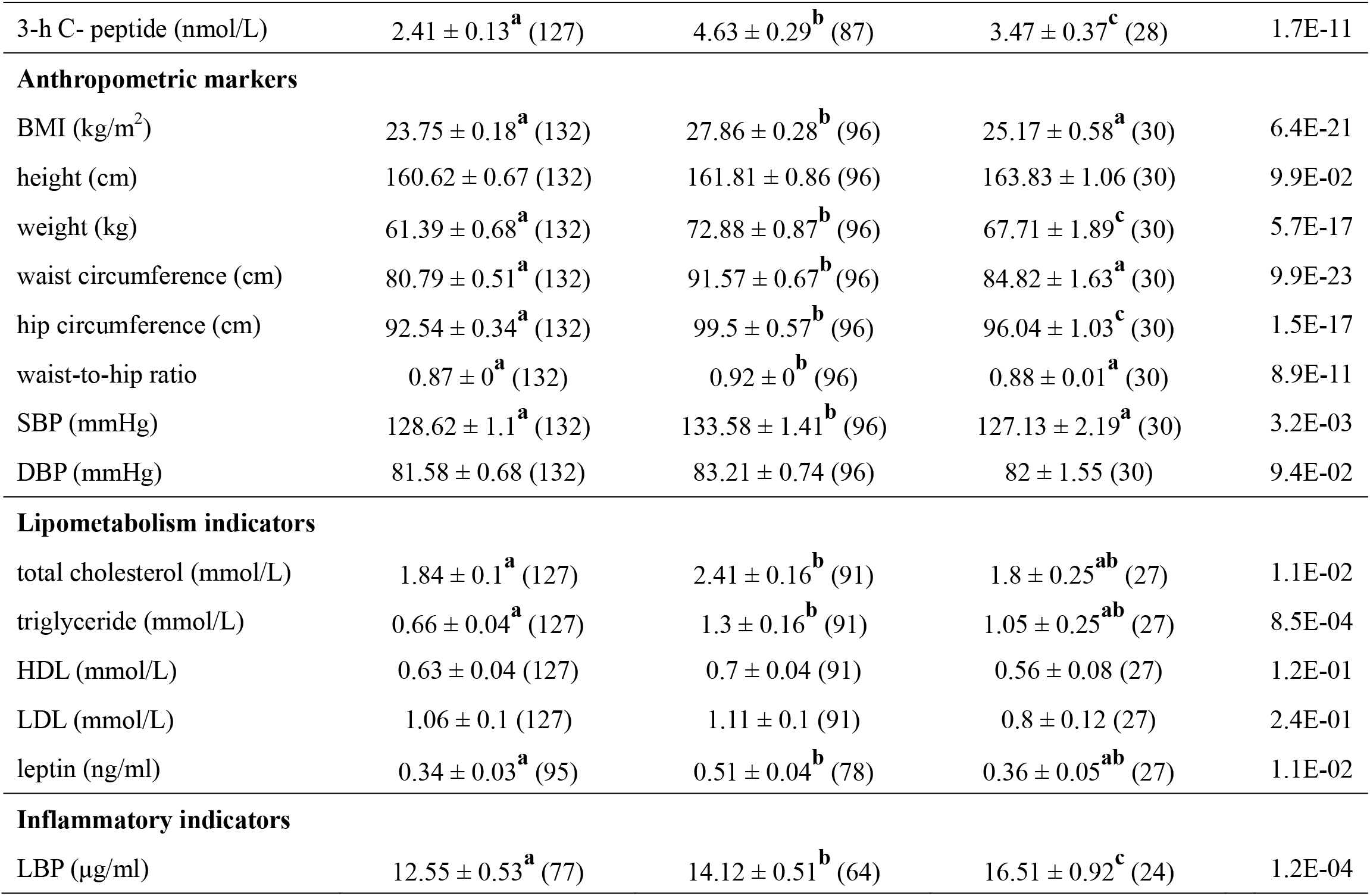
Comparisons of clinical parameters among unsupervised-stratification based clusters in discovery cohort.

Taken together, participants in Cluster 1 showed high insulin sensitivity and the lowest glucose levels and serum lipids. Participants in Cluster 2 had serious insulin resistance and high serum lipids levels. Participants in Cluster 3 had insulin deficiency, with the highest levels of blood glucose and chronic inflammation.

### **Differences in gut microbiota in unsupervised**-stratification based **clusters**

To investigate the differences in the gut microbiota among three unsupervised-stratification based clusters, we performed 16S rRNA gene V3-V4 region sequencing on fecal samples collected from all of the participants. Although there were no significant differences in diversity and richness of the gut microbiota (Fig. S2), the PCA of phylogenetic-ILR (PhILR, phylogenetic-isometric log ratio transformation)–transformed Euclidean distances and score plots of the linear discriminant analysis (LDA) showed that the structure and composition of the gut microbiota differed significantly among the three clusters (Fig. 2A and 2B). Moreover, the distance between Cluster 1 and Cluster 3 was larger than it was between Cluster 1 and Cluster 2 (Fig. 2C).

**FIG 2.**
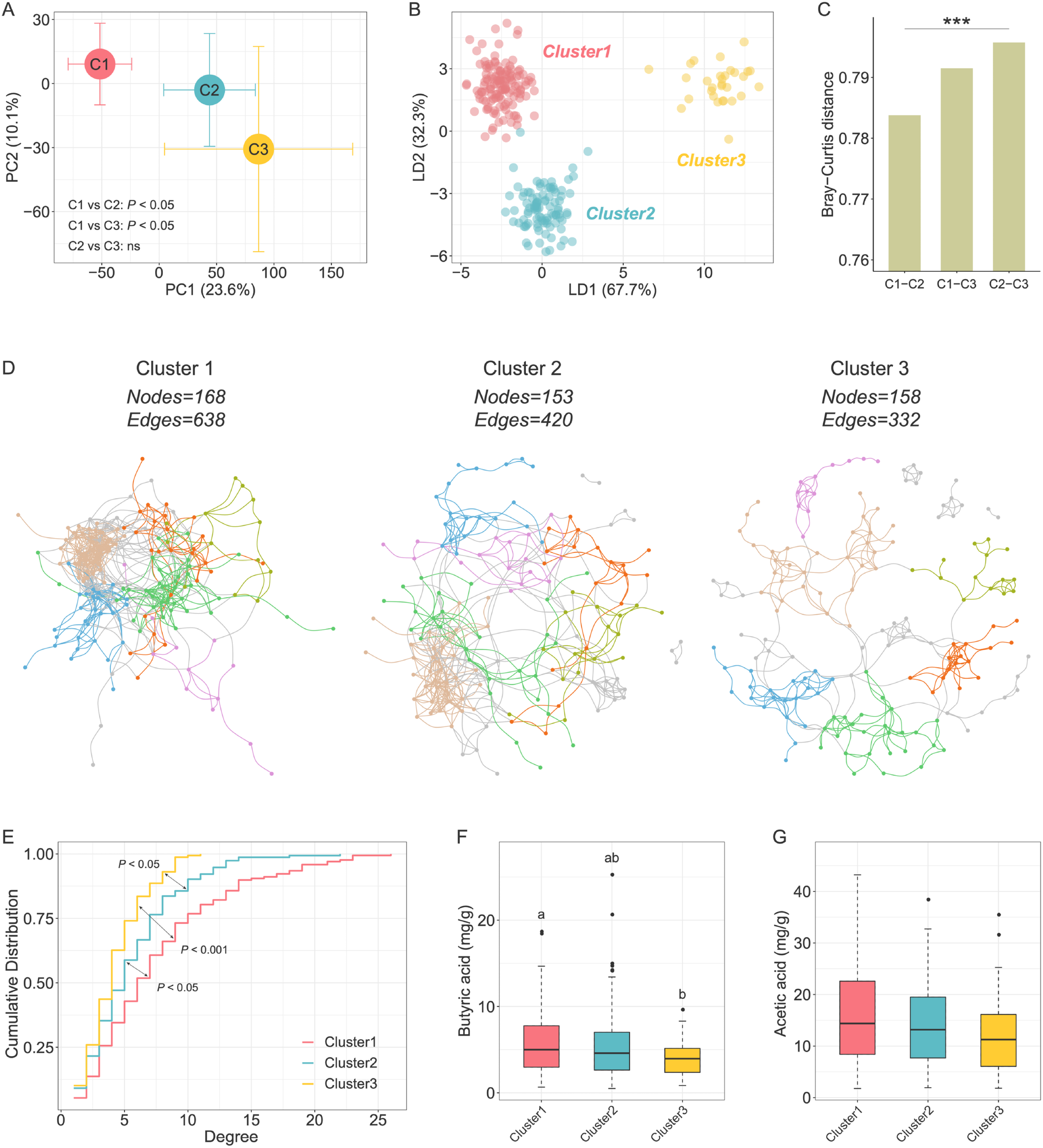
Characterization of the gut microbiota in unsupervised-stratification based clusters. **(A)** PCA of PhILR-transformed Euclidean-distance based on the abundance of ASVs. The circles and error bars indicate the mean and standard errors of the mean. The comparison of gut microbiota structures among three clusters were tested by permutational multivariate analysis of variances (PERMANOVA, permutations = 9,999). **(B)** LDA score plot of the gut microbiota structure of the three clusters. **(C)** Between-sample Bray-Curtis distances of the gut microbiota of three clusters. Kruskal-Wallis test, \*\*\**P* < 0.001. **(D)** Visualization of constructed networks based on Pearson correlation coefficient. The first six modules with a large number of nodes are shown in different colors, and the other modules are shown in grey. **(E)** The degree centralities of networks from three clusters. Kolmogorov-Smirnov tests were used to test the differences in cumulative distributions. Comparisons of **(F)** fecal butyric acid and **(G)** fecal acetic acid concentration among clusters. Boxes, whiskers, and outliers denote values as described for Fig. 1E. The Wilcoxon rank-sum test was used for comparisons between two clusters (adjusted by FDR). Clusters with common characters were not significantly different (FDR > 0.05).

Then we constructed a co-abundance network of prevalent ASVs (shared by more than 20% samples) in each cluster based on Pearson correlation coefficient to explore the ecological relationship of the members in gut microbial community (Fig. 2D). We calculated the topological parameters of networks in three clusters to explore whether any differences existed in complexity among the microbial networks. The total number of nodes was similar (168, 153, and 158) among the three networks, whereas the total number of edges varied relatively (638, 420, and 332), in particular, the network in Cluster 3 had the lowest number of edges. The network density, which is defined as the ratio of the number of actual edges and the number of possible edges, decreased progressively in the three clusters (the values of density in Clusters 1, 2, and 3 were 0.045, 0.036, and 0.027, respectively). Moreover, the network degree centrality, a measure of the relative connectivity of each node in a network, decreased from Cluster 1 to Cluster 3 (Fig. 2E). These results suggested that Cluster 1 had more microbial interactions than the other two clusters. As bacteria act as functional groups (guilds) in the gut ecosystem(23), we next clustered the 188 nodes (i.e., ASVs) of the three networks into 29 co-abundance groups (CAGs). Correlation analysis between CAGs and clinical parameters showed significant correlation between gut microbiota and glycometabolism, insulin secretion, and lipid levels (Fig. S3).

Moreover, we measured the content of short-chain fatty acids (SCFAs) in the fecal samples of all participants. SCFAs are important microbiota-derived metabolites and have been proved to be associated with glycometabolism. We found that Cluster 1 had the highest concentration of fecal butyric acid, whereas Cluster 3 had the lowest (Fig. 2F). The difference of acetic acid concentration among three clusters was similar with butyric acid (Fig. 2G). We further examined the genes involved in the production of butyric and acetic acid, e.g., *buk* for butyrate and *fhs* for acetate production. The gene abundances showed a pattern similar to that of the fecal butyric and acetic acid concentration among three clusters (Fig. S4).

Taken together, the β-diversity of gut microbiota, gut microbial network topology, and the capacity for producing butyrate and acetate were significantly different among these three clusters.

### **Features of gut microbiota in unsupervised**-stratification based **clusters**

To identify the cluster-specific microbiome features of three clusters, we compared the abundance of ASVs among the three clusters using Wilcoxon-rank sum test (FDR < 0.05, |log2-fold change| > 1). We found that 67 ASVs were significantly different between at least two groups (Fig. 3 top panel; see Table S1 in the supplemental material). Only 14 of the 67 ASVs showed significantly altered abundance between Clusters 1 and 2, and 36 ASVs were significantly altered between Clusters 1 and 3, whereas the abundance of 43 ASVs changed significantly between Clusters 2 and 3. β-diversity analysis based on the Bray-Curtis distance showed significant correlations between the profiles of 67 cluster-specific ASVs and all ASVs (Mantel test, R = 0.4, *P* = 0.001; Fig. S5, Procrustes Analysis, *P* < 0.001). In addition, the Mantel statistic based on Euclidean distance of 67 cluster-specific ASVs and Euclidean distance of 16 clinical variables used for unsupervised clustering, showed that the alteration pattern of microbiome features was significantly associated with the clinical phenotype (Mantel test, R = 0.14, *P* = 0.003).

**FIG 3.**
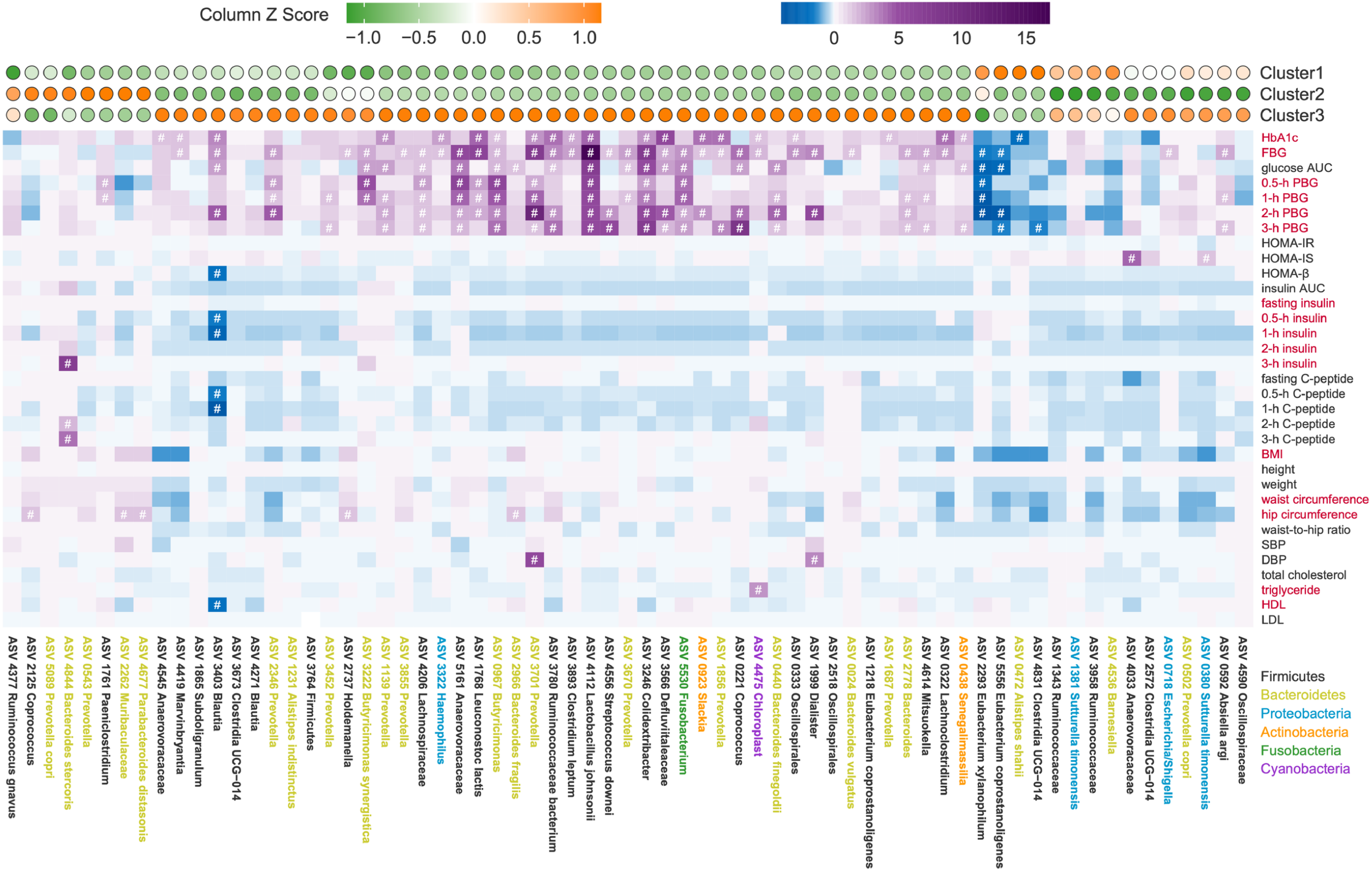
Cluster-specific microbiome features identified based on unsupervised-stratification based clusters. Top panel: The colors of circles indicate the scale-transformed mean abundance of the 67 ASVs in each cluster. These ASVs were clustered with a Spearman correlation coefficient and ward linkage based on their scale-transformed abundance values. Middle panel: Association between ASVs and clinical variables. The colors denote the correlation coefficients. *P*-values were adjusted by “BH”. # adjusted *P* < 0.25 was considered to be statistically significant based on the instruction of MaAslin2. Age and gender were considered to be covariates. Red text on the right indicates the variables used for classification. Bottom panel: Taxonomy of ASVs. Colors represent the phyla.

We subsequently assessed the correlation between the cluster-specific ASVs and all of the host clinical variables based on a modified general linear model (Fig. 3). Among the 67 cluster-specific ASVs, we found that 50 ASVs were correlated with at least one clinical variable. Four ASVs (three belonged to Clostridia: *Eubacterium xylanophilum* ASV2293, *Eubacterium coprostanoligenes* ASV5556, and *Clostridia UCG-014* ASV4831; one belonged to Bacteroidia: *Alistipes shahii* ASV0472), which were significantly higher in Cluster 1, were negatively correlated with the parameters related to glucose intolerance. Most of the ASVs (39 ASVs) enriched in Cluster 3 were significantly positively correlated with parameters related to glucose intolerance, but five of them were positively correlated with parameters related to lipid metabolism. Among them, two ASVs showed significant correlation with insulin-related variables. ASV3403, belonging to *Blautia*, was negatively correlated with HOMA-β, but ASV4033 enriched in Cluster 3, was positively correlated with HOMA-IS. One of the ASVs enriched in Cluster 2, *Bacteroides stercoris* ASV4844, was positively correlated with insulin and C-peptide levels. Other ASVs (*Paeniclostridium* ASV1761, *Coprococcus* ASV2125, *Muribaculaceae* ASV2262, and *Parabacteroides distasonis* ASV4677) enriched in Cluster 2 were positively correlated with parameters related to glucose intolerance or hip circumference. Taken together, the results indicated a significant correlation between cluster-specific microbiome features and clinical phenotypes.

### Stratification in testing cohort based on cluster-specific microbiome features

We next developed machine-learning classifiers based on a random forest algorithm by a leave-one-out cross-validation to distinguish individuals in one cluster from those in other two clusters using the 67 cluster-specific ASVs. Receiver operating characteristic curve analysis suggested that the models had high prediction power with area under the curve (AUC) ranging from 0.88 to 0.94 (Fig. 4A). Then we tested whether these features of gut microbiota could distinguish the subjects from different glycometabolism clusters in the testing cohort, who were the survey participants at the same community two years ago. We assigned the participants in the testing cohort to the nearest one of the three clusters based on Euclidean distance of the clinical variables that had been used for clustering in the discovery cohort except for the 3-h glucose level and 3-h insulin level (because these two variables were not available for the testing cohort) (Fig. 4B). The principal components analysis (PCA) based on clinical variables, which we used for classification, revealed a separation among the three clusters of testing cohort (Fig. 4B). HOMA-IR and HOMA-β were significantly higher in Cluster 2 than in the other two clusters (Fig. 4C and 4D), which was similar to the differences identified among the clusters in the discovery cohort. In addition, Cluster 1 showed the lowest levels of glucose AUC and Cluster 3 showed the highest levels (Table S2), which suggested that Cluster 1 had the best glucose metabolism. The gut microbiota structure of the three clusters in the testing cohort were clearly separated from each other (Fig. 4E) and the score plot of the LDA showed that the gut microbiota structure was different among clusters and not cohorts (Fig. S6). To explore whether the participants in the different clusters in the testing cohort could be distinguished by the microbial profile, we developed machine-learning classifiers again using the abundance of the 67 cluster-specific ASVs in the testing cohort. Receiver operating characteristic curve analysis suggested that the models had a moderate prediction power with an AUC ranging from 0.87 to 0.94 (Fig. 4F). The classification for the testing cohort suggested that the cluster-specific ASVs reflected the microbiota alterations associated with abnormal glycometabolism.

**FIG 4.**
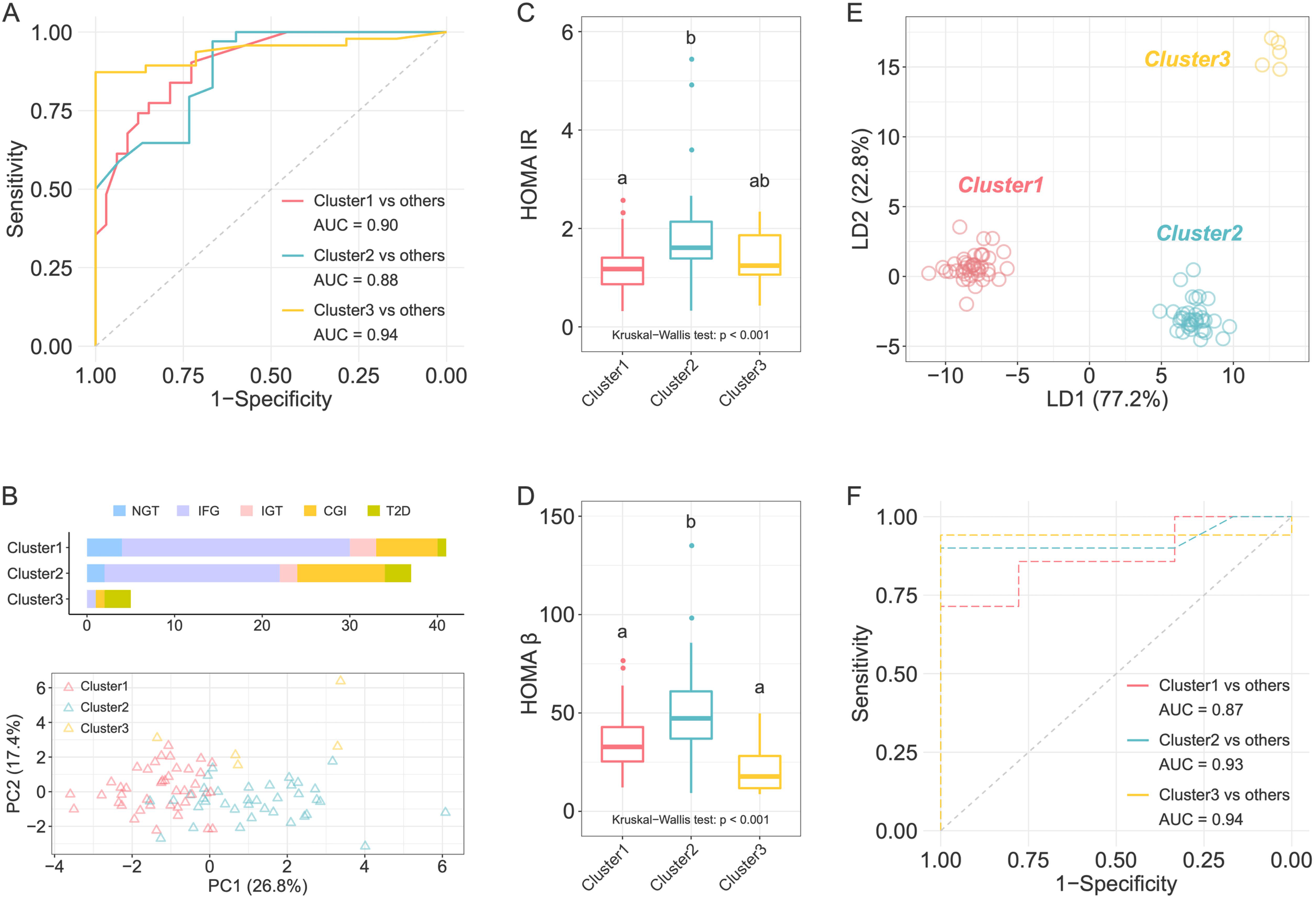
Classification based on cluster-specific microbiome features in the testing cohort. **(A)** Receiver-operating characteristic (ROC) curves for classification of individuals in one cluster from the other two clusters in the discovery cohort. The random forest classifier was constructed based on leave-one-out cross-validation using the 67 cluster-specific ASVs. **(B)** The number of participants in each cluster in the testing cohort, with colors indicating glycemic categories according to ADA criteria, and PCA plot showing the different clinical phenotype of three clusters. Comparisons **(C)** HOMA-IR and **(D)** HOMA-β among clusters. Kruskal-Wallis test *P*-value is shown at the bottom of each plot. Boxes, whiskers, and outliers denote values as described for Fig. 1E. The Wilcoxon rank-sum test was used for comparisons between two clusters (adjusted by FDR). Clusters with common characters were not significantly different (FDR > 0.05). **(E)** LDA score plot of the three clusters based on the abundance of ASVs. **(F)** ROC curves for classification of individuals in one cluster from the other two clusters in the testing cohort using the 67 cluster-specific microbial features identified in the discovery cohort.

## DISCUSSION

In the current population-based cross-sectional study, we showed that unsupervised stratification of patients with abnormal glycometabolism based on more inclusive clinical parameters could help identify microbiome features more robustly associated with glycometabolism. We confirmed the association between identified microbiome features and glycometabolism in a validation cohort.

The microbiome composition of individuals with T2D has been controversial among studies (24). *Bifidobacterium* and *Bacteroides* were the most reported genera containing microbes related with T2D. *Bifidobacterium* has been reported to be potentially protective against T2D in most studies, whereas only one study has reported conflicting result (2, 25–28). *Bacteroides* has been reported to be negatively correlated with T2D in five crosssectional studies and positively correlated with T2D in three studies that had involved some type of treatment (4, 5, 24, 29). One of the reasons for this unreliable relationship between gut microbiota and T2D is that individuals within the same blood glucose range are heterogeneous in insulin sensitivity and secretion as well as lipid metabolism, which are also associated with gut microbiota (21). In our study, based on 16 variables that combined blood glucose levels and parameters related to insulin resistance and dyslipidemia, we classified 258 individuals into three clusters with unique metabolic characteristics: Cluster 1 was characterized by the lowest blood glucose levels, insulin sensitivity, and lowest lipid levels; Cluster 2 was characterized by a moderate level of blood glucose, serious insulin resistance, and high levels of cholesterol and triglyceride; and Cluster 3 was characterized by the highest blood glucose levels and insulin deficiency. The Swedish All New Diabetics study reported that clusters identified based on more inclusive indexes and an unsupervised method showed different risk of diabetic complications (13). This result suggested that the stratification based on more inclusive clinical parameters was better than that based only on blood glucose levels, because it not only could separate people within different glucose levels and insulin levels but also could predict the risk of diabetic complications. Based on the unsupervised-stratification based clusters, we identified cluster-specific microbiome features that not only were related to glycometabolism but also were available for population classification in another general cohort. This finding implied an association between these cluster-specific microbiome features and glycometabolism. Thus, our research findings suggested that the stratification combining the blood glucose levels and indicators related to insulin resistance and dyslipidemia could make it possible to identify the microbiome features associated with abnormal glycometabolism.

The identified cluster-specific microbiome features may contribute to the progression of abnormal glycometabolism. For example, we found that two ASVs belonging to *Prevotella copri* were enriched in Clusters 2 and 3, respectively, and one ASV belonging to *Bacteroides vulgatus* was enriched in Cluster 3. Both Cluster 2 and Cluster 3 were characterized by the most resistance to insulin. One study found that *P. copri* and *B. vulgatus* were the strongest driver species for the positive association between HOMA-IR and microbial branched-chain amino acids (BCAAs) biosynthesis in Danish individuals without diabetes and further found that *P. copri* caused insulin resistance and impaired glucose intolerance by changing the circulating serum levels of BCAAs in mice (30). Transplantation of *B. vulgatus* resulted in insulin resistance in recipient mice (31). Therefore, these two bacteria may have contributed to a high level of insulin resistance in Clusters 2 and 3 in our work. Furthermore, we also found a high abundance of *Ruminococcus gnavus* ASV4377 in Clusters 2 and 3. Studies reported that *R. gnavus* is a mucin-degrading bacterium that may directly break the integrity of gut barrier and is associated with inflammatory bowel diseases (32–35). The disruption of gut barrier may lead to the translocation of endotoxins produced by gut bacteria to the host. In our study, the level of LBP, a load marker of gut-derived antigens, was higher in Clusters 2 and 3, which suggested an increased level of plasma endotoxin load produced by gut bacteria. The increased circulating endotoxin load would induce chronic inflammation, which is a driving factor for insulin resistance and dyslipidemia (36). Thus, the disordered gut microbiome, such as increased levels of *R. gnavus*, may contribute to the insulin resistance and dyslipidemia by disrupting gut barrier, elevating circulated endotoxin load, and inducing chronic inflammation. Moreover, Cluster 1 enriched the ASVs belonging to *Barnesiella*. Some species of *Barnesiella* have been reported to produce acetate (37). In addition, high levels of acetate and butyrate concentration, as well as the functional genes involved in the production of these metabolites were observed in Cluster 1. Studies have revealed that acetate suppresses body fat accumulation and inflammation in obese or diabetic rodents through multiple mechanisms (38–41). Butyrate also has been shown to improve gut integrity by increasing the tight junction assembly (42), inducing mucin synthesis (43), and decreasing gut bacterial transport across the epithelium (44). Thus, the individuals in Cluster 1 may have benefited from the integrity of the intestinal barrier, which was protected by higher acetate/butyrate concentration, and may have avoided the elevated endotoxin load that can be induced by chronic inflammation. Taken together, the cluster-specific microbiome features found in the current study may have contributed to the distinct glycometabolism phenotype of the three clusters. The contribution and mechanism of these bacteria in the progression of glycometabolism disorder need to be further experimentally verified.

In this study, we showed that unsupervised stratification based on blood glucose, insulin and lipid levels led to the identification of cluster-specific gut microbiome features associated with glycometabolism. A well-stratified cohort is a prerequisite for the identification of bacteria associated with glycometabolism. Upon further validation in larger cohorts and follow-up with a longer duration, the elucidation of the mechanism of these identified cluster-specific microbiome features in the progression of abnormal glycometabolism may lead to the future development of biomarkers for early diagnosis and therapeutic treatment.

## MATERIALS AND METHODS

### Ethical approval

The protocols for both studies were approved by the by the Human Research Ethics Committee of Shanghai General Hospital (2009KY037, 2013KY083) before the procedure of enrollment. This clinical trial was registered in the Chinese Clinical Trial Registry under number ChiCTR-IPC-14005346. All of the participants signed an informed consent form before sample collection.

### Overview of the cohorts

Patients at the Sijing Community Health Service Center of Songjiang District participated in a survey about type 2 diabetes (T2D). We recruited 267 individuals from a diabetes survey taken in 2014, which was considered to be the discovery cohort in the analysis. To test our findings in the discovery cohort, we recruited 86 individuals from a diabetes survey taken in 2012 as the testing cohort.

The participants were asked to fast overnight (more than 10 h) to collect the fasting venous blood. After physical examination and fasting venous blood collection, we performed a 3-h OGTT (75 g glucose) and collected venous blood samples at 30, 60, 120, and 180 min. The blood samples were set at room temperature for 30 min and then centrifuged to obtain the serum. The serum of fasting venous blood was divided into two parts: one was used to evaluate the fasting blood glucose, blood lipid, and inflammation; and the other was immediately stored at −80°C for quantification of LBP and leptin. Stool samples were collected on the day of the physical examination and stored at −80°C quickly until fecal DNA extraction.

### Biochemical assays

The levels of HbA1c, serum glucose, serum insulin, serum C-peptide, triglyceride, total cholesterol, HDL cholesterol, and LDL cholesterol were determined at Shanghai General Hospital, Shanghai Jiao Tong University School of Medicine. Enzyme-linked immunosorbent assays (ELISAs) were used to quantify the levels of LBP (Hycult Biotech, PA, USA) and leptin (DL develop, Wuhan, China) in the lab of Shanghai Jiao Tong University.

### Unsupervised-stratification

We used 16 clinical variables (HbA1c, five-time-point blood glucose levels and insulin levels during OGTT, BMI, waist circumference, hip circumference, triglyceride, HDL) to stratify the discover cohort. We performed unsupervised-stratification in an R environment (version 3.6.1). Euclidean distances were calculated (the *vegdist* function in the “vegan” package) for the standardized clinical variables (scaled to a mean of 0 and S.D. of 1) to complete the clustering analysis. Individuals with outlier variables (absolute standardized levels ≥ 5) were excluded from the clustering analysis. We performed the K-Mediods clustering algorithm using the *pam* function in the “cluster” package to complete the clustering procedure. The silhouette-width was calculated using the *silhouette* function in the “cluster” package. Clusterboot algorithm from the “fpc” package was used to assess the stability of clusters. Finally, other than nine participants who had at least one outlier variable, 258 participants were clustered into three clusters.

We used the median values of the 14 selected clinical variables in the discovery cohort, except for the 3-h glucose and 3-h insulin levels (because these two variables were not available for the testing cohort), to assign participants to clusters in the testing cohort. We took the nearest neighbors of the three cluster centers based on Euclidean distances. After we removed three participants with outlier variables, 83 participants were used in the testing cohort.

### Statistical analysis of clinical data

Statistical analysis of clinical data was performed in an R environment (version 3.6.1). The difference in the clinical variables among clusters was tested by Kruskal-Wallis test. For differences between two clusters, Wilcoxon rank-sum test was used (adjusted by FDR). FDR values were converted into a character-based display in which common characters represented clusters that were not significantly different (the *multcompLetters* function in the “multcompView” package). The Pearson chi-square test was performed to compare the differences in categorical data. *P* < 0.05 (for Kruskal-Wallis test) and FDR < 0.05 (for Wilcoxon rank-sum test) were considered to have a significant difference.

### Fecal DNA extraction and 16S rRNA gene V3-V4 region sequencing

We extracted fecal microbial DNA based on the previously published method (45). A total of 353 samples were sequenced in four batches on Miseq system (Illumina, San Diego, CA, USA). The sequencing library of 16S rRNA gene V3-V4 regions was prepared as previously described (46), according to a modified version of the manufacturer’s instructions.

We used QIIME2 software (v2018.11) (47) to process and analyze the 16S rRNA gene read pairs. The raw sequence data were demultiplexed, denoised, and filtered for chimeric reads with the DADA2 plugin (48) to obtain the frequency table and representative sequence file of amplicon sequence variants (ASVs). After we removed the ASVs considered to be contaminants (49), the decontamination table composed of 353 sample and 5,448 ASVs was downsized to 10,000,000 to standardize sequence depth. We used the representative sequence file for taxonomic annotation using the SILVA database (version 138).

### Functional gene prediction

The functional genes for producing butyrate (*but*, butyrate kinase) and acetate (*fhs*, formate-tetrahydrofolate ligase) were predicted based on 16S rRNA gene information by using Phylogenetic Investigation of Communities by Reconstruction of Unobserved States (PICRUSt2) (50).

### Bioinformatics and statistical analysis of microbiota data

The following analyses of microbiota were performed by R (version 3.6.1). The α-diversity of each sample was calculated with Shannon index, Simpson index, Observed ASVs, and Faith’s phylogenetic diversity (PD whole tree) (R-packages “picante”, “phyloseq”, and “ape”). Structural differences in gut microbiota were assessed by β-diversity based on PhILR-transformed Euclidean distance using the R-packages “phyloseq” and “philr”. Random forest model was trained to distinguish the individuals in one cluster from those in other two clusters using the *train* function in the R-package “caret”.

Wilcoxon rank-sum test (adjusted by FDR) was used to analyze the difference of α-diversity index and identify the significantly different ASVs between two clusters. ASVs were considered to be significantly different between two clusters when the FDR was lower than 0.05 and the absolute value of the logarithmic (base 2) fold change (|log2-fold changeļ) in relative abundance was greater than 1. Permutational multivariate analysis of variance (PERMANOVA; permutations = 9,999) was used to assess the structural difference between different clusters.

### Network construction

In each cluster, prevalent ASVs shared by more than 20% of the samples were used to construct the microbial association network. Networks of Clusters 1, 2, and 3 were generated based on Pearson correlations of the prevalent ASVs using the R-package “WGCNA”. The correlations with *P* < 0.05 (adjusted by BH) were retained for further analysis. The layout of nodes and edges was determined by the Fruchterman-Reingold layout algorithm using the correlation efficient as weight. The topological characteristics calculation and visualization of the networks were performed using the R-package “igraph”. Kolmogorov-Smirnov test (the *ks.test* function in the “stats” package) was used to compare the network’s topological characteristics between clusters. *P* < 0.05 was considered to have a significant difference.

Next, the ASVs from the three networks were clustered using the “ward.D2” (the hclust function in the “vegan” package) based on the correlation distance which were converted from correlation values. Permutational MANOVA (permutations = 9999, P < 0.01) was used to determine whether the two clades of the cluster tree were not significantly different and clustered into one CAGs.

### Association of microbiome features and clinical phenotypes

To calculate the associations between microbiome features and clinical phenotypes, we used a modified general linear model as implemented by MaAsLin2 (Multivariate microbial Association with Linear Models), which combines an arcsine square root transformed analysis of relative abundances in a standard multivariable linear model while adjusting for gender and age. The *P* values were adjusted by Hochberg-Benjamini procedure. According to the instructions of the “Maaslin2” package, adjusted *P-*value lower than 0.25 was considered to be significant.

### Quantification of fecal short chain fatty acids (SCFAs)

Fecal SCFAs were quantified by gas chromatography/mass spectrometry (GC/MS) as previously described (2). Wilcoxon rank-sum test was used to analyze the difference of SCFAs between clusters.

### Data availability

The raw sequence data reported in this paper have been deposited (PRJCA013291) in the Genome Sequence Archive (GSA) database under accession number CRA008952.

## ACKNOWLEDGMENTS

Funding to C. Z. was supported by grants from the National Key Research and Development Project (2019YFA0905600) and the National Natural Science Foundation of China (31922003 and 81871091). Funding to X. D. was supported by grants from the National Natural Science Foundation of China (81870594) and Clinical research plan of SHDC (No.SHDC2020CR1016B).

## SUPPLEMENTAL MATERIAL

Supplemental material is available online only.

**FIG S1 The variations of HOMA-β among members within each ADA group.** The colors indicate different ADA groups, the size of each circle denotes the value of HOMA-β in each individual. (PDF file, 21 KB).

**FIG S2 Violin plots of the differences in α-diversity of gut microbiomes among three clusters.** The shaded background for each cluster reflects the data distribution, boxes show the interquartile ranges (IQRs), the point in each box represents the median, the whiskers denote the lowest and highest values that were within 1.5 times the IQR from the first and third quartiles, and the outliers are shown as individual points. Wilcoxon rank-sum test was used to compare the difference between two clusters. * *P* < 0.05. (PDF file, 358 KB)

**FIG S3 Associations between clinical parameters and gut microbiota.** Left panel: The colors of circles indicate the scale-transformed mean abundance of the CAGs in each cluster. Right panel: Association between CAGs and clinical variables. The colors denote the correlation coefficients. The CAGs were clustered with a Spearman correlation coefficient and ward linkage based on their correlation coefficients with clinical parameters. P-values were adjusted by “BH”. + adjusted *P* < 0.25 was considered to be statistically significant based on the instruction of MaAslin2. Age and gender were considered to be covariates. (PDF file, 263 KB)

**FIG S4 Relative abundance of genes involved in bacterial butyric and acetic acid production.** Comparisons of **(A)** butyric production gene, *buk*, butyrate kinase; **(B)** acetic acid production gene, *fhs*, formate-tetrahydrofolate ligase. Boxes show the medians and the interquartile ranges (IQRs), the whiskers denote the lowest and highest values that were within 1.5 times the IQR from the first and third quartiles, and the outliers are shown as individual points. Wilcoxon rank-sum test was used to compare the difference between two clusters. (PDF file, 214 KB)

**FIG S5 Similar β-diversity pattern between all ASVs and 67 cluster-specific ASVs.** Procrustes analysis and Mantel test are used to compare the structural difference of all ASVs and 67 cluster-specific ASVs based on Bray-Curtis distance. Procrustes analysis, *P* < 0.001. Mantel test, R = 0.4, *P* = 0.001. (PDF file, 35 KB)

**FIG S6 LDA score plot of the gut microbiota structure of clusters in discovery cohort and testing cohort.** The colors denote the different clusters. The solid and hollow circles denote the individuals in discovery cohort and testing cohort, respectively. (PDF file, 360 KB)

**Table S1 Cluster-specific microbiomes features identified in discovery cohort.** Wilcoxon rank-sum test (adjusted by FDR) was used to compare the abundances of ASVs between two clusters. 67 ASVs with FDR < 0.05 and ļlog2-fold changeļ > 1 were considered as cluster-specific microbiome features. (XLSX file, 18 KB)

**Table S2 Comparisons of clinical parameters among clusters in testing cohort.**

Data are shown as mean ± sem (num). A Kruskal-Wallis test was used to compare the differences among the three clusters. For comparisons between two clusters, we used the Wilcoxon rank-sum test (adjusted by FDR) for continuous variables and used the chisquared test for categorical variables. Data with common characters were not significantly different (FDR > 0.05). The full name of abbreviations, computational formula of HOMA-IR, HOMA-IS, and HOMA-β as described for Table 1. (DOCX file, 26 KB)

